# A universal model for carbon dioxide uptake by plants

**DOI:** 10.1101/040246

**Authors:** Han Wang, I. Colin Prentice, William K. Cornwell, Trevor F. Keenan, Tyler W. Davis, Ian J. Wright, Bradley J. Evans, Changhui Peng

## Abstract

The rate of carbon uptake by land plants depends on the ratio of leaf-internal to ambient carbon dioxide partial pressures^1^, here termed *χ*. This quantity is a key determinant of both primary production and transpiration and the relationship between them. But current models for *χ* are empirical and incomplete, contributing to the many uncertainties afflicting model estimates and future projections of terrestrial carbon uptake^2^,^3^. Here we show that a simple evolutionary optimality hypothesis^4^,^5^ generates functional relationships between *χ* and growth temperature, vapour pressure deficit and elevation that are precisely and quantitatively consistent with empirical *χ* values from a worldwide data set containing > 3500 stable carbon isotope measurements. A single global equation embodying these relationships then unifies the empirical light use efficiency model with the standard model of C_3_ photosynthesis^1^, and successfully predicts gross primary production as measured at flux sites. This achievement is notable because of the equation′s simplicity (with just two parameters, both independently estimated) and applicability across biomes and plant functional types. Thereby it provides a theoretical underpinning, grounded in eco-evolutionary principles, for large-scale analysis of the CO_2_ and water exchanges between atmosphere and land.

## Main

Current Earth System Models (ESMs) disagree even on the most basic processes in the global carbon cycle, including terrestrial CO_2_ uptake^2^,^3^ – suggesting a need to revisit foundational questions in ecosystem science^7^,^8^. Depending on their history and purpose, ESMs represent plant CO_2_ uptake either with the standard model of Farquhar et al.^1^, which accurately describes the instantaneous environmental and physiological controls of photosynthesis, or with the empirical light use efficiency (LUE) model, which can predict primary production over weeks to months^6^,^9^. These approaches have served for the past three decades as parallel frameworks for relating primary production to environmental drivers, but the connection between them remains tenuous^9^. Moreover, large-scale implementations of both require independent information to be provided, such as photosynthetic capacities (*V*_*cmax*_ and *J*_*max*_) and the ratio of leaf-internal (*c*_*i*_) to ambient (*c*_*a*_) CO_2_ concentrations (here termed *χ*) in the Farquhar model, and response functions for various environmental factors in the LUE model. There is no accepted general way to do this^10^,^11^, and as a result, different implementations of apparently the same model can give very different answers in different ESMs.

The biochemical reactions of photosynthesis are critically dependent on the value of *χ*^12^ CO_2_ diffuses into leaves through the stomata (microscopic pores in the leaf surface) towards the chloroplasts, where reducing power derived from solar energy is used to assimilate CO_2_ into organic forms through the Calvin cycle. *χ* is tightly regulated by the responses of stomatal aperture to environment. *χ* determines the availability of CO_2_ for assimilation, and thus constrains both the carboxylation‐ and electron transport-limited photosynthetic rates. However, current models that explicitly predict *χ* represent only its response to moisture, and even this is represented by several approximate and non-equivalent formulations (for more information on the theoretical background see Supplementary Methods S1)^13^. A firm basis for the prediction of *χ* is thus an essential step towards a first-principles representation of terrestrial plant carbon uptake. Here we derive a theory for the dependencies of *χ* on growing-season air temperature (*T*_*g*_), vapour pressure deficit VPD (*D*_*g*_), and elevation (*z*) based on the least-cost hypothesis^4^,^5^, which states that plants minimize the combined costs of maintaining the capacities for carboxylation (maintaining the activity of Rubisco, the primary carboxylating enzyme, and other photosynthetic proteins) and transpiration (maintaining living tissues to support water transport) required to achieve a given assimilation rate. The theory is tested against effective growing-season values of *χ* derived from a global compilation of stable carbon isotope (δ^13^C) measurements on leaves of C_3_ plants (Fig. S1). The additional hypothesis of co-limitation between carboxylation‐ and electron transport-limited photosynthetic rates is then used to provide a universal model of gross primary production (GPP), which unifies the Farquhar and LUE models.

Logit transformation of the predicted optimal value of *χ* (termed *χ*_*∘*_) yields remarkably simple theoretical partial relationships with each of the three environmental predictor variables (Supplementary Methods S2). The predicted effects of each variable are shown to be quantitatively consistent with those inferred from the data, within their uncertainties (Fig. 1, Table 1). Theory and data agree that logit (*χ*) rises by ~ 0.0545 per degree due to both increased assimilation costs (the affinity of Rubisco for CO_2_ *versus* O_2_ declines at higher temperatures) and reduced water transport costs (the viscosity of water also declines); falls by 0.5 per unit increase of ln *D*_*g*_ due to the increase in transpiration costs imposed by increasing *D*; and falls by ~ 0.0815 per km elevation due to reduced O_2_ pressure (increasing the affinity of Rubisco for CO_2_) and increased transpiration costs (because the saturated vapour pressure of water remains constant while the actual vapour pressure, *ceteris paribus,* declines). Thus, *χ* increases with temperature by ~ 0.01 K^−1^, decreases with VPD by ~ 0.1 kPa^−1^, and decreases with elevation by ~ 0.01 km^−1^. By imposing the theoretical values for the three environmental effects on *χ*, we estimate an intercept of 1.189, close to the fitted value of 1.168. The fully linearized theoretical model is then:

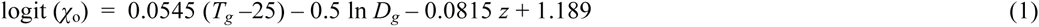

which is statistically indistinguishable from the fitted model for *χ* (Table 1).

**Figure 1.**
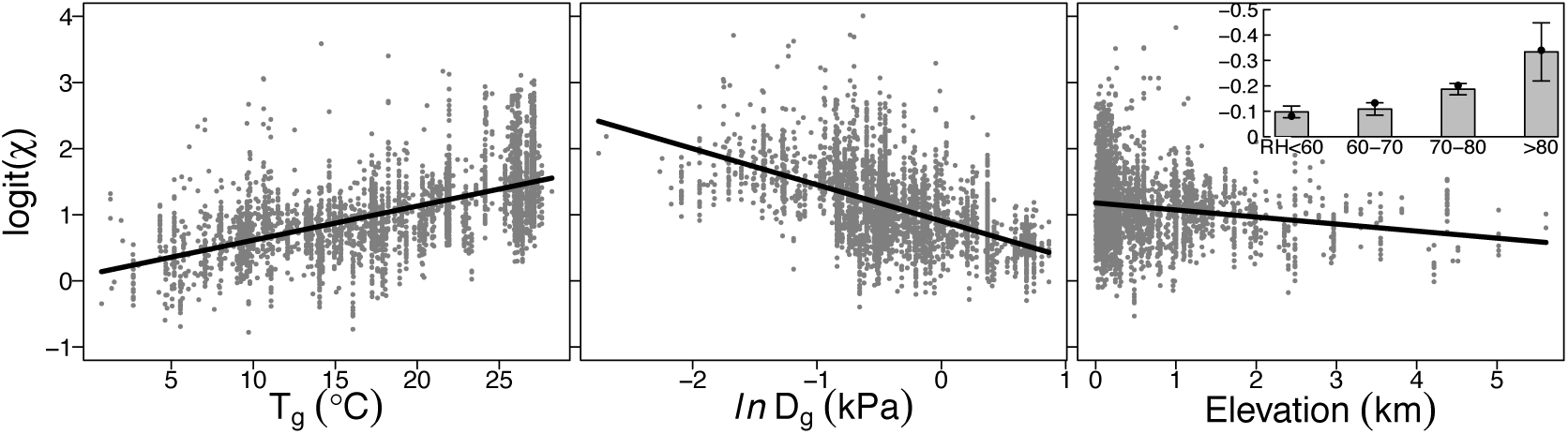
Partial residual plots from the regression of logit-transformed *χ* against environmental predictors. *χ*, the ratio of leaf-internal to ambient CO_2_ partial pressures. ΔT_*g*_, = growing-season mean temperature *T*_*g*_ - 25°C. ln *D*_*g*_, natural logarithm of growing-season mean vapour pressure deficit. Inset shows elevation responses for relative humidity (RH, %) classes, compared to predicted responses (black dots) evaluated at the centre of each class.

**Table 1.** Regression summaries. Logit-transformed values of the ratio of leaf-internal to ambient CO_2_ partial pressure (*χ*), derived from δ^13^C measurements, were regressed against the difference between growing-season mean temperature *T*_*g*_ and 25°C (*ΔT*_*g*_, °C), the natural logarithm of growing-season mean VPD (ln *D*_*g*_, kPa), and elevation (*z*, km). Theoretical values are partial derivatives with respect to each predictor, evaluated for standard conditions (*T*_*g*_ = 25 °C, *D*_*g*_ = 1 kPa, *z* = 0 km).

| Predictor | Fitted coefficient | Confidence intervals |  | Theoretical value | Model R<sup>2</sup> |
| --- | --- | --- | --- | --- | --- |
|  |  | 2.5% | 97.5% |  |  |
| $\Delta T_g$ | 0.0515 | 0.0456 | 0.0575 | **0.0545** | 0.391 |
| $\ln D_g$ | -0.5478 | -0.6111 | -0.4846 | **-0.5** | |
| $z$ | -0.1065 | -0.1315 | -0.0815 | **-0.0815** | |
| intercept | 1.1680 | 1.0464 | 1.2896 | **1.189** |  |

Equation (1) yields *χ*_*∘*_ = 0.77 under standard conditions (*T*_*g*_ = 25 °C, *D*_*g*_ = 1 kPa, *z* = 0 km). The predicted elevation effect increases with relative humidity (RH), becoming arbitrarily large as RH approaches 100% (Supplementary Methods S2). As predicted, the fitted (negative) slope of logit (*χ*) with elevation becomes larger with RH, most steeply at high RH (Fig. 1). Using an independent dataset of instantaneous CO_2_ and water exchange measurements^14^, we also show - consistent with equation (1) - that the single parameter determining the sensitivity of *χ*_*∘*_ to VPD is influenced by temperature, but not by VPD (Table S1).

*χ*_*∘*_ values from equation (1) are consistent with observed *χ* across biomes (r = 0.51) (Fig. 2). Highest values are in hot, wet, low-elevation sites (tropical forests), lowest in cold and/or dry and/or high-elevation sites (deserts, polar and alpine vegetation). *χ*_*∘*_ ranges globally from 0.4 to almost 1.0 (Fig. S2). The reduction from the equator towards mid-latitudes is due to aridity while that in high latitudes is due to declining temperatures (Fig. S3). The elevation effect on *χ* is long-known, but has not been satisfactorily explained^15^,^16^. By predicting it in the same framework that accounts for climate effects, we have resolved a long-standing conundrum, showing that the unit cost of photosynthesis is reduced while that of transpiration is increased with elevation, leading to reduced *χ*_*∘*_.

**Figure 2.**
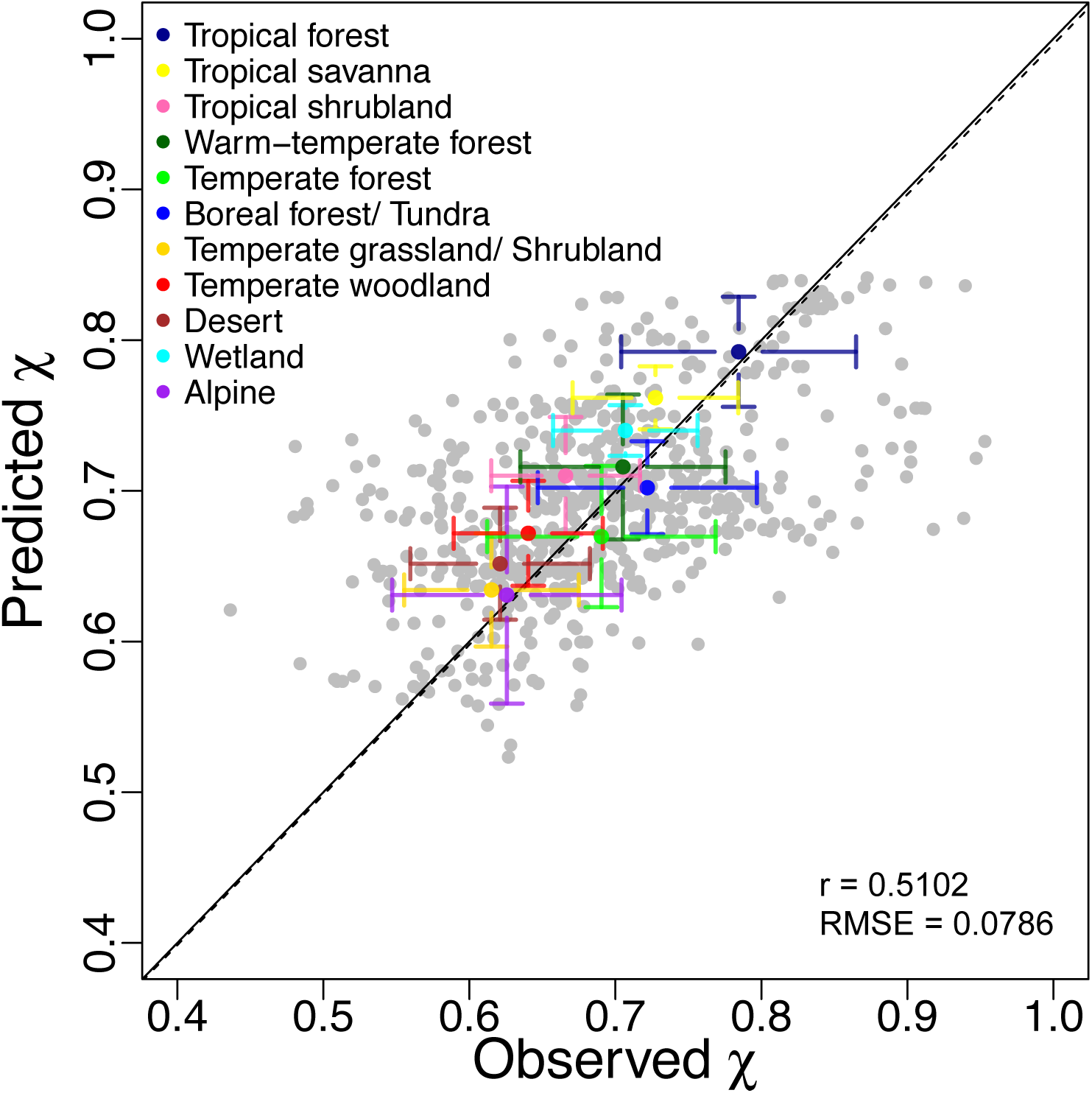
Site-mean leaf-internal to ambient CO_2_ partial pressures (χ). Predictions from the theoretical model driven by three environmental variables (Table 1); observations from the global δ^13^C dataset. Mean and standard deviation are shown for each biome. Biome type for each site were assigned based on mega-biome classification from BIOME430 for consistency of definitions and wetland and alpine types from literatures records. The regression line through the origin is imposed as the black solid line; the dashed line is the 1:1 line.

Some published analyses focused on leaf δ^13^C as a palaeoclimate indicator^15,17,18^. Unlike the well-documented effect of aridity on δ^13^C, these analyses detected no temperature effect, even through it is predicted both by the earlier Cowan-Farquhar criterion^19^ and by the least-cost hypothesis^4^ and has been shown both in short-term experiments^14^,^20^ and in field data^4^. Mean annual precipitation (MAP) has previously been used to represent plant water availability; the lack of an observed temperature effect might then be an artefact, because MAP tends to increase with temperature. We showed a significant (but much weakened) effect of temperature when MAP was substituted for VPD (Table S2) based on our much larger data set. However the controlling variable is VPD, not MAP.

No significant difference was found between woody and non-woody plants (Fig. S4). The most parsimonious interpretation for the statistical significance of plant functional type (PFT) differences in *χ* detected here and elsewhere is as an indirect effect caused by different PFTs’ climatic preferences^14^. This interpretation is strongly supported by Fig. S4, which shows that differences in ^13^C discrimination among PFTs are predicted correctly by the universal model. We did however show a slightly lower *χ* for evergreen needleleaf trees than the other PFTs (Fig. S4). This is consistent with higher intrinsic water use efficiency in conifer forests than broadleaf forests, and could be attributed to the lower permeability of gymnosperm wood (the consequence of narrower conducting elements)^21^. According to our analysis, the estimated water cost is 20% higher in gymnosperms, even though the resulting difference in *χ* from this component is slight (3%: Supplementary Methods S2). The slightly overestimated *χ* for evergreen needleleaf trees by the universal model, and the spread of observed *χ* values around the central tendency, suggest that distinguishing hydraulic influences on the controls of unit transpiration costs, in particular, might further improve predictability.

We detected a significant negative response of *χ* to soil pH, explaining an additional 5% of variance. This finding is consistent with a soil-calcium restoration experiment that enhanced annual evapotranspiration by 20%^22^, and other findings of high *χ* on acid substrates^23^. The framework could be extended to consider N uptake costs, which may be higher (favouring investment in water transport) on less fertile soils.

The co-limitation hypothesis, stating that the two photosynthetic processes of carboxylation and transport are coupled such that photosynthetic rates limited by those two processes are equal under typical daytime conditions, provides the necessary next step towards a universal model of GPP^24^,^25^. The hypothesis implies adjustment of *V*_*cmax*_ in time and space to match environmental conditions^25^. Extensive field measurements also point to an optimal maximum rate of electron transport, *J*_*max*_ that maximizes the photosynthetic benefits minus the costs of maintaining the electron-transport chain^26^ (Supplementary Methods S4). We can thereby eliminate both *V*_*cmax*_ and *J*_*max*_ as independent predictors, to derive a first-principles model:

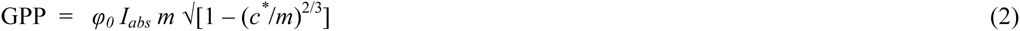

where

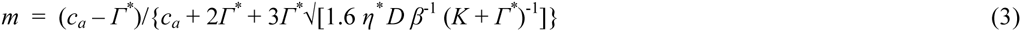

Here *φ*_*0*_ is the intrinsic quantum yield of photosynthesis (1.02 g C mol^−1^)^27^, *I*_*abs*_ is the absorbed photosynthetic photon flux density (PPFD, mol m^−2^ s^−1^), *Γ*^*^ is the photorespiratory compensation point (Pa), *n*^*^ is the viscosity of water relative to its value at 25°C, *β* represents the ratio of carboxylation and transpiration costs at 25°C (*β* ≈ 240, estimated from the constant in equation 1), and *c*^*^ is the unit carbon cost for the maintenance of electron transport capacity, ≈ 0.41 (estimated from observed *J*_*ma*_*x-V*_*cmax*_ ratios) (Fig. S5). LUE is the product of *φ*_*0*_, *m* and the square-root term in equation (2); thus, GPP is proportional to *I*_*abs*_, which can be calculated from incident PPFD and remotely sensed green vegetation cover. Predicted monthly GPP compared well with monthly GPP derived from CO_2_ flux measurements (Fig. 3). Predicted global total annual GPP is 120 Pg C, within the accepted range^28^.

**Figure 3.**
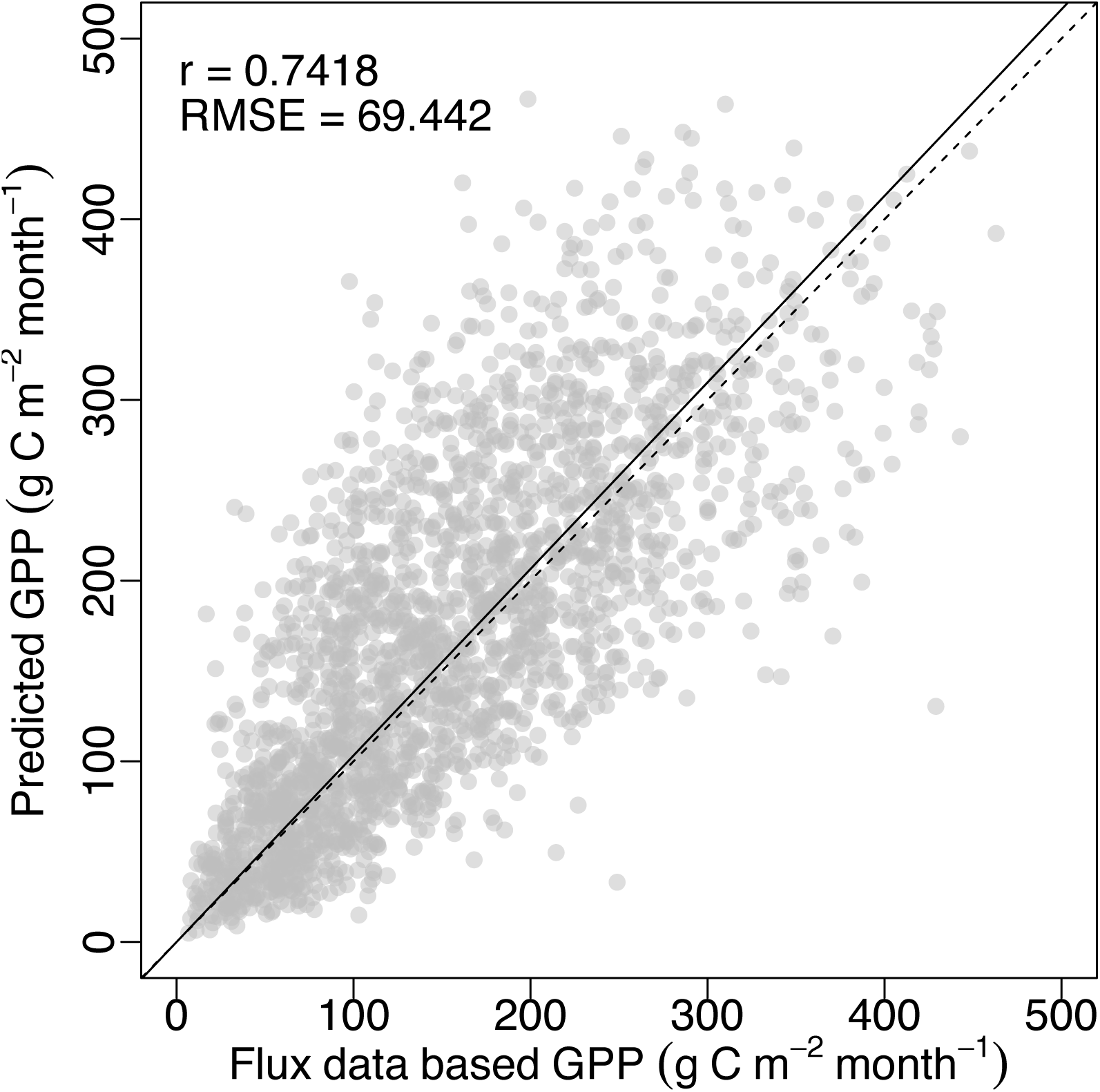
Monthly gross primary production (GPP). Predictions from equations (2) and (3); observations based on CO_2_ flux data in the FLUXNET archive. The regression line through the origin is imposed as the black solid line; the dashed line is the 1:1 line.

Additional testable predictions arise, for example on the controls of net primary production (NPP). Our results intriguingly parallel findings from metabolic scaling theory, whereby monthly NPP was predicted and found to be proportional to growing-season length and biomass but to show a weakly negative response to temperature^29^. Many potential complications, such as the environmental dependencies of mesophyll conductance^20^, the influence of soil fertility factors on nutrient acquisition costs, and the differences among photosynthetic pathways, have been neglected so far; yet this simplistic model’s predictive skill suggests a promising route to an improved predictive understanding of terrestrial carbon and water cycling.

## Acknowledgements

Research supported by an Australian Research Council Discovery grant (‘Next-generation vegetation model based on functional traits’) to ICP and IJW, a National Basic Research Programme of China (2013CB956602) grant to HW and CP, an Australian National Data Service grant (‘Ecosystem production in space and time’) to ICP, and Terrestrial Ecosystem Research Council (TERN) grants (‘Ecosystem Modelling and Scaling Infrastructure’) to ICP and BJE. TERN and ANDS are supported by the Australian Government National Collaborative Infrastructure Strategy (NCRIS). TFK acknowledges support from a Macquarie University Research Fellowship. We thank Yan-Shih Lin, Vincent Maire, Belinda Medlyn and Beni Stocker for discussions. The paper is a contribution to the AXA Chair Programme on Biosphere and Climate Impacts and Imperial College’s initiative on Grand Challenges in Ecosystems and the Environment. In addition to authors of this paper, data were provided by Margaret Barbour, Lucas Cernusak, Todd Dawson, David Ellsworth, Graham Farquhar, Howard Griffiths, Claudia Keitel, Alexander Knohl, Peter Reich, Dave Williams, Radika Bhaskar, Hans Cornelissen, Anna Richards, Susanne Schmidt, Fernando Valladares, Christian Korner, Ernst-Detlef Schulze, Nina Buchmann and Lou Santiago. We used ‘free and fair use’ eddy-covariance data acquired by the FLUXNET community and, in particular, by the following networks: AmeriFlux (US Department of Energy, Biological and Environmental Research, Terrestrial Carbon Program (DE-FG02-04ER63917 and DE-FG02-04ER63911)), AsiaFlux, CarboEuropeIP, Fluxnet-Canada (supported by CFCAS, NSERC, BIOCAP, Environment Canada, and NRCan), OzFlux and TCOS-Siberia. We acknowledge the financial support to the eddy-covariance data harmonization provided by CarboEuropeIP, FAO‐ GTOS-TCO, iLEAPS, Max Planck Institute for Biogeochemistry, National Science Foundation, University of Tuscia, Universite Laval and Environment Canada and US Department of Energy and the database development and technical support from Berkeley Water Center, Lawrence Berkeley National Laboratory, Microsoft Research eScience, Oak Ridge National Laboratory, University of California-Berkeley, University of Virginia.

## Methods

### Theory for the environmental controls on *χ*

According to the least-cost hypothesis^4^, optimal *χ* minimizes the combined costs of maintaining the capacities for carboxylation and transpiration:

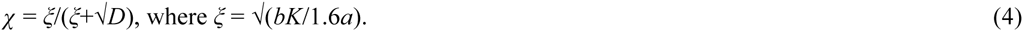

The ratio of xylem repiration to transpiration capacity (*a*) depends *inter alia* on the viscosity of water; the ratio of mitochondrial respiration to carboxylation capacity (*b*) is generally taken as constant^1^. *D* is the vapour pressure deficit (VPD); *K* is the effective Michaelis-Menten coefficient of Rubisco.

Logit transformation of (4) yields:

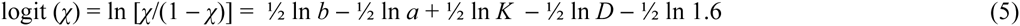

Temperature affects *χ* through *a* via viscosity, and *K via* the Michaelis-Menten coefficients for carboxylation and oxygenation. Elevation affects *χ* through *K* and *D via* the partial pressures of oxygen and water vapour respectively. Separating environmental effects from invariant quantities in (5) leads to the definition of a constant term:

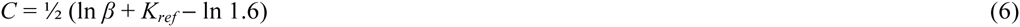

where *K*_*ref*_ and *β* are the values of *K* and the ratio *b/a,* respectively, under standard conditions (T = 298 K, *z* = 0).

Using the Vogel equation for viscosity^31^, the Arrhenius equation for biochemical rate parameters and the barometric formula relating atmospheric pressure to elevation, we evaluated the partial derivatives of χ with respect to T, *D* and *z* at *T* = 298 K, z = 0 and *D* = 1 kPa. *C* was estimated as the intercept in a Generalized Linear Model (GLM) fitted to the data with imposed regression coefficients (the calculated partial derivatives) for all three environmental effects (Supplementary Methods S2).

### Testing the theory with global δ^13^C data

Vascular-plant leaf stable carbon isotope data were compiled from published and unpublished sources by Cornwell *et al.* (submitted). The data can be downloaded from Dryad (Data link http://datadryad.org/review?doi=doi:10.5061/dryad.3jh61). Inferred carbon isotope discrimination (Δ) values for 3549 leaf samples of C3 plants^32^ (Supplementary Methods S3) were converted to estimates of *χ* by a standard method. The Climatic Research Unit CL2.0 10-minute gridded monthly climatology^33^ of mean, maximum and minimum temperatures and relative humidity provided mean temperature (*T*_*g*_, °C) and vapour pressure deficit (*D*_*g*_, kPa) values for the period with daily mean temperatures > 0°C. Logit (*χ*) values were entered in a GLM with Δ*T*_*g*_ = *T*_*g*_‐ 25°C, ln *D*_*g*_, and site-specific elevation (*z*, km) as predictors. Standard errors estimated by the GLM were combined quadratically with standard errors for the uncertainty of the Rubisco discrimination parameter b’, the latter obtained by generating 10^4^ normally distributed values of b’ (mean = 27, standard deviation = 0.27) and repeating the estimation of *χ* and the GLM fitting 10^4^ times with different *b’* values.

### Light-use efficiency model

The model proposed by Wang et al.^8^ assumed that the electron-transport and Rubisco-limited rates of photosynthesis (*A*_*J*_, *A*_*c*_) are co-limiting under typical daytime conditions^24 25 34^, allowing GPP to be predicted from A_J_. LUE is the product of φ_0_ and the CO_2_ limitation term (denoted here by *m*) in the model of ref. 8. Incorporating the exact equation for *χ*_*∘*_ (equation 8 in ref. 4) yields:

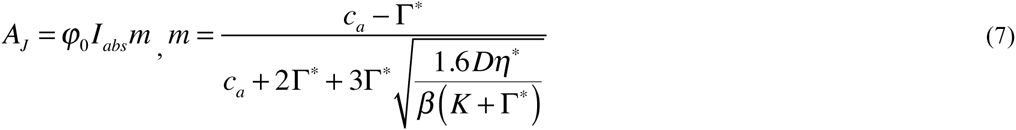

However, (7) implies that the light response of *A*_*J*_ is linear up to the co-limitation point, i.e. the maximum electron-transport rate (*J*_*max*_) is arbitrarily large. In reality *J*_*max*_ limitation can be be significant, especially at high temperatures. We therefore modify (7) to consider a non-rectangular hyperbola relationship between *A*_*J*_ and *I*_*abs*_^35,36^:

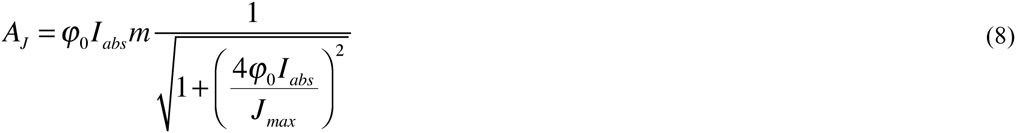

We further assume that there is a cost associated with *J*_*max*_ equal to the product of *J*_*max*_ and a constant (*c*^*^), and that optimal *J*_*max*_ maximizes the benefit (*A*_*J*_) minus this cost. The optimal ratio *J*_*max*_/*V*_*cmax*_ at the growth temperature is then:

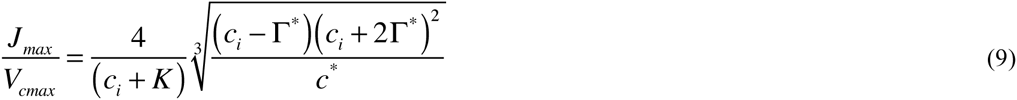

where *c*_*i*_ = *χc*_*a*_, and *c*^*\**^ can be estimated from data in ref. 26. We evaluated (9) at each grid cell in the CRU CL1.0 climatology^37^ and regressed the results against *T*_*g*_ (Fig. S5), indicating a strong relationship consistent with observations^26^. The LUE model is accordingly revised to (Supplementary Methods S4):

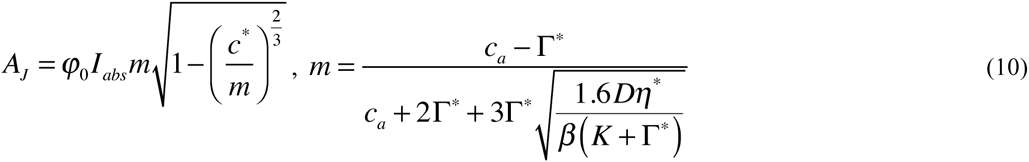

### GPP data-model comparison

Equations (2)-(3) yielded modelled site-specific monthly GPP values for comparison with values independently derived from eddy-covariance measurements of CO2 exchange in the Free and Fair Use subset of the FLUXNET archive, using a consistent gap-filling procedure (Supplementary Methods S5). For the modelled values, monthly LUE was estimated based on temperature and vapour pressure extracted from CRU time-series (TS 3.22) data at 0.5°C resolution^38^ and site-observed *c*_*a*_. Monthly absorbed PPFD was estimated as the product of PPFD (0.45 times the WATCH incident surface shortwave radiation^39^, divided by 0.22 J μmol^−1^) and the MODIS Enhanced Vegetation Index (EVI), equated to the fraction of photosynthetically active radiation absorbed by foliage^40^. To match the WATCH data resolution, EVI was upscaled to the 0.5° grid cell in which each site was located.

## Author contributions

The least-cost theory was first proposed by I.J.W. and further developed by I.C.P. I.C.P. and H.W. derived the predictions. H.W. carried out all the analyses and constructed the Figures and Tables. W.K.C. originated and compiled the Δ^13^C data set. B.J.E., T.W.D. and I.C.P. developed and tested the flux partitioning method; T.W.D. developed the global flux database and all the GPP computations. T.F.K contributed on quantifying the regression uncertainties. H.W. and I.C.P. wrote the first draft; all authors contributed to the final draft.

## References

1. Farquhar, G. D., von Caemmerer, S., & Berry, J. A., A biochemical model of photosynthetic CO_2_ assimilation in leaves of C_3_ species. Planta 149, 78–90 (1980).

2. Ciais, P., et al. Carbon and other biogeochemical cycles. In: Climate Change 2013: the Physical Science Basis. Contribution of Working Group I to the Fifth Assessment Report of the Intergovernmental Panel on Climate Change. 465–570 (Cambridge University Press, 2014).

3. Friedlingstein, P., et al. Uncertainties in CMIP5 climate projections due to carbon cycle feedbacks. Journal of Climate 27, 511–526 (2014).

4. Prentice, I. C., Dong, N., Gleason, S. M., Maire, V., & Wright, I. J., Balancing the costs of carbon gain and water transport: testing a new theoretical framework for plant functional ecology. Ecology letters 17, 82–91 (2014).

5. Wright, I. J., Reich, P. B., & Westoby, M. Least-cost input mixtures of water and nitrogen for photosynthesis. The American Naturalist 161, 98–111 (2003).

6. Monteith, J. L., Solar radiation and productivity in tropical ecosystems. Journal of Applied Ecology 9, 747–766 (1972).

7. Prentice, I. C., Liang, X., Medlyn, B. E., & Wang, Y. P., Reliable, robust and realistic: the three R's of next-generation land-surface modelling. Atmospheric Chemistry and Physics 15, 59876005 (2015).

8. Wang, H., Prentice, I. C., & Davis, T. W., Biophsyical constraints on gross primary production by the terrestrial biosphere. Biogeosciences 11, 5987–6001 (2014).

9. Medlyn, B. E., Physiological basis of the light use efficiency model. Tree Physiology 18, 167176 (1998).

10. Ali, A. et al. A global scale mechanistic model of the photosynthetic capacity. Geoscientific Model Development Discussions 8, 6217–6266 (2015).

11. Cai, W. et al. Large differences in terrestrial vegetation production derived from satellite-based light use efficiency models. Remote Sensing 6, 8945–8965 (2014).

12. De Kauwe, M. G., et al. Forest water use and water use efficiency at elevated CO_2_: a model-data intercomparison at two contrasting temperate forest FACE sites. Global Change Biology 19, 1759–1779 (2013).

13. Medlyn, B. E., et al. Reconciling the optimal and empirical approaches to modelling stomatal conductance. Global Change Biology 17, 2134–2144 (2011).

14. Lin, Y.-S. et al. Optimal stomatal behaviour around the world. Nature Climate Change 5, 459–464 (2015).

15. Korner, C., Farquhar, G. D., & Roksandic, Z. A global survey of carbon isotope discrimination in plants from high altitude. Oecologia 74, 623–632 (1988).

16. Friend, A., Woodward, F. & Switsur, V., Field measurements of photosynthesis, stomatal conductance, leaf nitrogen and 8^13^C along altitudinal gradients in Scotland. Functional Ecology 3, 117–122 (1989).

17. Diefendorf, A. F., Mueller, K. E., Wing, S. L., Koch, P. L., & Freeman, K. H., Global patterns in leaf ^13^C discrimination and implications for studies of past and future climate. Proceedings of the National Academy of Sciences 107, 5738–5743 (2010).

18. Kohn, M. J., Carbon isotope compositions of terrestrial C_3_ plants as indicators of (paleo) ecology and (paleo) climate. Proceedings of the National Academy of Sciences 107, 19691–19695 (2010).

19. Cowan, I. R., & Farquhar, G. D., Stomatal function in relation to leaf metabolism and environment. Symposia of the Society for Experimental Biology 31, 471–505 (1977).

20. Caemmerer, S. & Evans, J. R., Temperature responses of mesophyll conductance differ greatly between species. Plant, Cell & Environment 38, 629–637 (2015).

21. Frank, D. C., et al. Water-use efficiency and transpiration across European forests during the Anthropocene. Nature Climate Change 5, 579–583 (2015).

22. Green, M. B., et al. Decreased water flowing from a forest amended with calcium silicate. Proceedings of the National Academy of Sciences 110, 5999–6003 (2013).

23. Maire, V., et al. Global effects of soil and climate on leaf photosynthetic traits and rates. Global Ecology and Biogeography 24, 706–717 (2015).

24. Maire, V., et al. The coordination of leaf photosynthesis links C and N fluxes in C_3_ plant species. PloS One 7, e38345 (2012).

25. Haxeltine, A. & Prentice, I. C., A general model for the light-use efficiency of primary production. Functional Ecology 10, 551–561 (1996).

26. Kattge, J. & Knorr, W., Temperature acclimation in a biochemical model of photosynthesis: a reanalysis of data from 36 species. Plant, Cell & Environment 30, 1176–1190 (2007).

27. Long, S. P., Postl, W. F., & Bolhar-Nordenkampf, H. R., Quantum yields for uptake of carbon dioxide in C3 vascular plants of contrasting habitats and taxonomic groupings. Planta 189, 226–234 (1993).

28. Beer, C., et al. Terrestrial gross carbon dioxide uptake: global distribution and covariation with climate. Science 329, 834–838 (2010).

29. Michaletz, S. T., Cheng, D., Kerkhoff, A. J., & Enquist, B. J., Convergence of terrestrial plant production across global climate gradients. Nature 512, 39–43 (2014).

30. Kaplan, J. O., Geophysical applications of vegetation modeling. (Doctoral Thesis, Lund University, 2001).

## References

31. Vogel, H., Temperaturabhangigkeitsgesetz der Viskositat von Flussigkeiten. Physik Z 22, 645-646 (1921).

32. Farquhar, G. D., Ehleringer, J. R., & Hubick, K. T., Carbon isotope discrimination and photosynthesis. Annual Review of Plant Biology 40, 503–537 (1989).

33. New, M., Lister, D., Hulme, M. & Makin, I., A high-resolution data set of surface climate over global land areas. Climate Research 21, 1–25 (2002).

34. Chen, J.-L., Reynolds, J. F., Harley, P. C., & Tenhunen, J. D., Coordination theory of leaf nitrogen distribution in a canopy. Oecologia 93, 63–69 (1993).

35. Smith, E. L., The influence of light and carbon dioxide on photosynthesis. The Journal of general physiology 20, 807–830 (1937).

36. Harley, P. C., Thomas, R. B., Reynolds, J. F., & Strain, B. R., Modelling photosynthesis of cotton grown in elevated CO_2_. Plant, Cell & Environment 15, 271–282 (1992).

37. New, M., Hulme, M. & Jones, P., Representing twentieth-century space-time climate variability. Part I: Development of a 1961–90 mean monthly terrestrial climatology. Journal of Climate 12, 829–856 (1999).

38. Harris, I., Jones, P., Osborn, T. & Lister, D., Updated high-resolution grids of monthly climatic observations-the CRU TS3.10 Dataset. International Journal of Climatology 34, 623–642 (2014).

39. Weedon, G. P., et al. The WFDEI meteorological forcing data set: WATCH Forcing Data methodology applied to ERA-Interim reanalysis data. Water Resources Research 50, 7505–7514 (2014).

40. Xiao, X., Zhang, Q., Hollinger, D., Aber, J. & Moore, B. I., Modeling gross primary production of an evergreen needleleaf forest using MODIS and climate data. Ecological Applications 15, 954–969 (2005).

